# Synergistic interaction of hyposalinity stress with *Vibrio* infection causes mass mortalities in oysters by inducing host microflora imbalance and immune dysregulation

**DOI:** 10.1101/2021.08.31.458472

**Authors:** Xin Li, Ben Yang, Chenyu Shi, Hebing Wang, Qi Li, Shikai Liu

**Affiliations:** Key Laboratory of Mariculture (Ocean University of China), Ministry of Education and College of Fisheries, Ocean University of China, Qingdao 266003, China; Laboratory for Marine Fisheries Science and Food Production Processes, Qingdao National Laboratory for Marine Science and Technology, Qingdao 266237, China

**Keywords:** Oyster, Digestive microbiota, Immunity, Hyposalinity stress

## Abstract

A sudden drop in salinity following extreme precipitation events usually causes mass mortalities of oysters exposed to pathogens in ocean environment. While how hyposalinity stress interacts with pathogens to cause mass mortality remains obscure. In this study, we performed an experiment by mimicking hyposalinity stress and pathogen infection with *V. alginolyticus* to investigate their synergistic effect on the mortality of infected oysters toward understanding of the interaction among environment, host, and pathogen. We showed that hyposalinity stress (10‰, 20‰ versus 30‰) did not significantly affect proliferation and virulence of *V. alginolyticus*, but significantly altered microbial composition and diversity in infected oysters. Microbial community profiling by 16S rRNA sequencing revealed disrupted homeostasis of digestive bacterial microbiota with increased abundance of several pathogenic bacteria, which may affect the pathogenesis in oysters. Transcriptome profiling of infected oysters revealed that a large number of genes associated with apoptosis and inflammation were significantly induced under hyposalinity, suggesting that hyposalinity stress may have triggered immune dysregulation in infected oysters. This work provides significant information in decoding mechanisms of synergistic interaction among environment factors, host genetics, and digestive microbiota, and how they contribute to pathogenesis.

**IMPORTANCE:** Revealing the response of oyster host and microbial community to the interference of multiple environmental factors is an important aspect of deciphering the complex pathogenic mechanism in oysters. We evaluated the synergistic effects of hyposalinity stress and *Vibrio alginolyticus* infection in oysters. Results showed that hyposalinity stress significantly caused mass mortalities of infected oysters by destroying digestive microbial community structure, and triggering excessive immune response in oysters. This work provides valuable information for deciphering the mechanisms of synergistic interaction among environmental factors, host, and pathogens, and how they contribute to pathogenesis.

## INTRODUCTION

Due to global warming, ice caps and glaciers melt rapidly (1), extreme precipitation events are projected to occur more frequently (2, 3), which poses increasing threats to marine ecosystems of coasts and estuaries (4–6), leading to the biodiversity of marine environments being extremely changed (7). As an important marine shellfish species, the oysters inhabit in estuaries and intertidal areas. Drastic changes in the salinity of estuaries and intertidal areas caused by heavy rainfall and inflow from rivers have a great effect on growth and reproduction of oysters (8). The fluctuations of salinity can cause physiological stress, promote pathogenesis of microbial infection, and aggravate disease outbreaks (9–11). As a benthic filter-feeders, oysters face tremendous exposure to pathogenic microbes, such as *Ostreid herpesvirus 1* (*OsHV-1*), *V. parahaemolyticus*, *V. harveyi* and *V. alginolyticus* (12–15). Disease outbreaks caused by pathogenic microbes pose a major threat to marine ecosystem and shellfish aquaculture in many countries.

Different *Vibrio* species produce a series of extracellular products, which are potential pathogenic factors for marine animals. These toxins mainly include extracellular hemolysin, protease, lipase, siderophore, exopolysaccharide and effectors delivered via the type III secretion systems (16–20). Salinity can regulate the progression of diseases by affecting pathogens, hosts, or host-pathogen interactions. It has been reported that salinity may play a role in *OsHV-1* transmission in oysters (21, 22). Understanding the interaction among environment, host, and pathogens is essential for exploring the complex aetiology of oyster disease caused by abiotic and biotic stress.

Alteration of the gut microbiota is involved in animal pathology (23, 24). Studies have shown that the dynamic changes of the microbiota are closely related to the development, health, metabolism and immunity of the host (25, 26). The digestive flora of many aquatic organisms not only promotes the body’s nutrient absorption by secreting digestive enzymes, but also protects the body against the invasion of pathogens (27). However, studies have also shown that the change of host’s external environmental conditions or pathogen invasion may cause the imbalance of host’s internal microbial homeostasis, thereby, symbiotic bacteria may transform into opportunistic pathogens to mediate the development of host diseases. For instance, opportunistic bacteria are involved in secondary bacterial infections in oysters that are immune-compromised due to viral infection (28). Therefore, the stability of the microbial community is critical to health and viability of the host (29).

High throughput sequencing has been applied to profile the microbial community and host transcriptome expression (28, 30). High throughput RNA sequencing (RNA-Seq) has been widely used to analyze various traits in oysters, including heat stress (31, 32), salinity stress (33, 34), and viral infections (35). These transcriptome analyses revealed differentially expressed genes related to various immune pathways, including macrophage (36), interleukin-1 receptor-associated kinase (37), C1q complement system (38), apoptosis pathway (39), and the NF-κB, toll-like receptor and MAPK signaling pathways (40). Among which, inflammatory factors and apoptosis play an important role in innate immune response. Pro-inflammatory mediators and oxidative stress can induce cell apoptosis (41–43). The apoptosis or necrotic cell death may also cause inflammation, however, if the inflammatory response is excessively severe, there is danger of continuous tissue damage, forming an auto-amplification loop that causes organ damage (42, 44). Studies have shown that sharp changes in salinity, when combined with additional co-occurring stressors, appeared to trigger dysregulation of the immune response in aquatic organisms, potentially making more susceptible to infectious diseases (45).

The risk of disease outbreaks in the marine environment depends on the interaction among the host, pathogens and environmental factors (46). However, most studies have investigated the host response to single environmental factor, making it difficult to characterize diseases with complex etiology. Thus, understanding of diseases caused by changes in the microbial community, the interaction among the host, environment and pathogens remains obscure. In the present work, we performed an experiment by mimicking hyposalinity stress and pathogen infection with *V. alginolyticus* to investigate their synergistic effect on the mortality of the infected oysters. We tested the effects of salinity on the growth and virulence of *V. alginolyticus*, and also investigated the transcriptome responses of infected oysters and digestive bacterial microbiota dynamics. The results showed that high mortality rate of infected oysters under hyposalinity stress was caused by the destruction of the bacterial balance and immune homeostasis, which suppressed the host resistance to *V. alginolyticus*. Investigation on the interactions among environment, host and pathogen provides insights into the complex mechanism of pathogenesis in oysters exposed to biotic and abiotic stressors. This work provides valuable information in decoding mechanisms of synergistic interaction among environment factors, host genetics, and digestive microbiota, and how they contribute to pathogenesis.

## RESULTS

### Oyster survival analysis

The mortality of oysters infected with *V. alginolyticus* at normal salinity (30‰) and hyposalinity (10‰) conditions was monitored throughout the disease progression (Fig. 1). Through a 12-day experiment, the infected oysters at normal salinity (30‰) showed 33% mortality, while oysters at hyposalinity (10‰) showed 100% mortality. No mortality was observed in the oysters injected with artificial seawater. The observation suggested that hyposalinity stress had a significant effect on the pathogenesis of *V. alginolyticus* to cause mortality of the oysters. All experimental procedures performed on oyster were conducted in conformity with the Management Rule of Laboratory Animals (Chinese Order No. 676 of the State Council, revised 1 March 2017). Protocols were approved by the Committee on the Ethics of Animal Experiments of Ocean University of China.

**Fig. 1.**
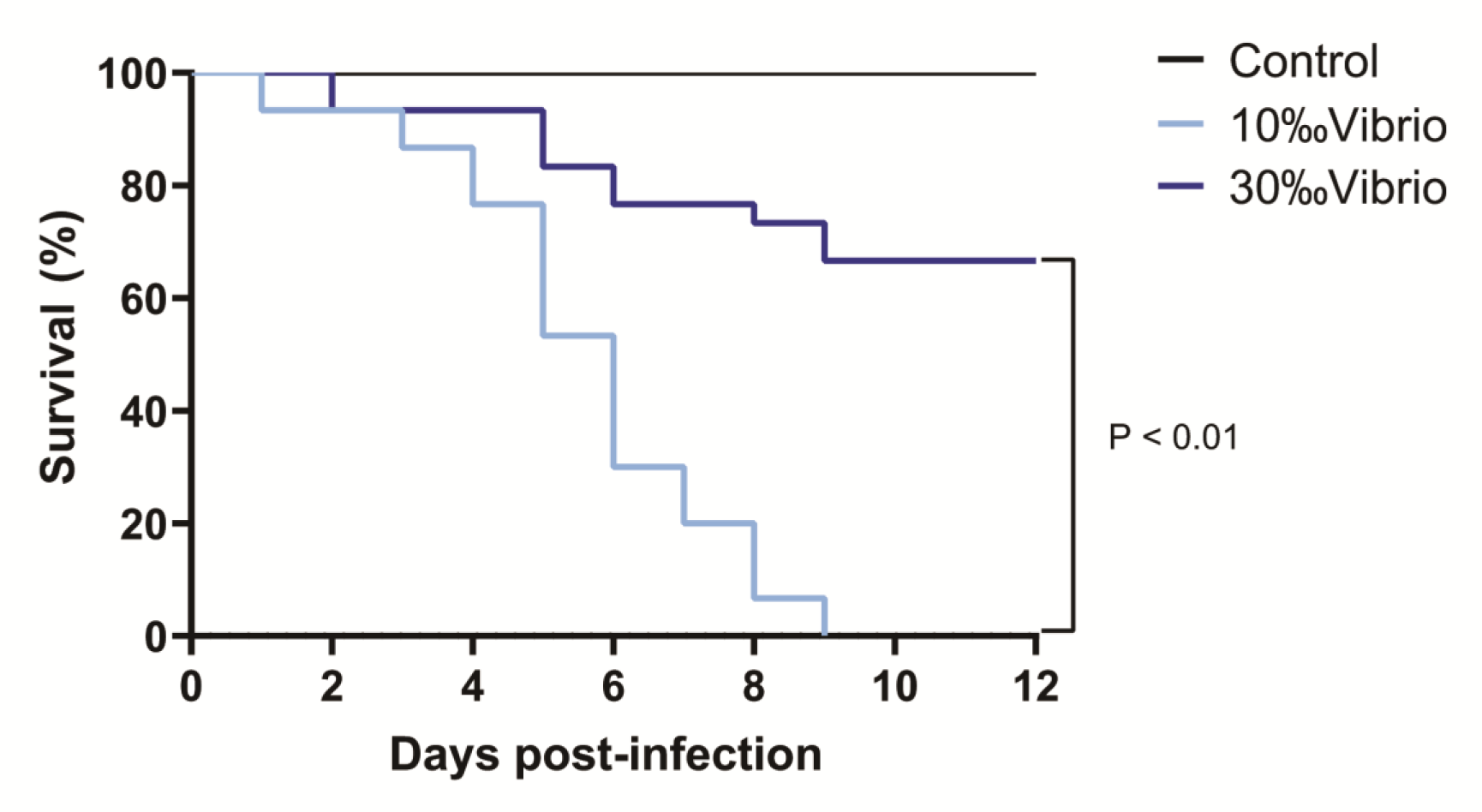
Survival analysis of *C. gigas* infected with *V. alginolyticus* at different levels of salinity. The mortality of oysters in each salinity group (n=90, in triplicate) was monitored for 12 days post infection with *V. alginolyticus*. ***, P value < 0.0001, log rank test; n = 30.

### Impact of hyposalinity on the growth and virulence of *V. alginolyticus*

To explore the effect of hyposalinity on *V. alginolyticus* infection in the oyster, we examined the growth and virulence of *V. alginolyticus* under different levels of salinity (Fig. 2). No significant difference was observed for growth of *V. alginolyticus* cultured with different levels of salinity (10‰, 20‰ and 30‰) after 12h (Fig. 2A). Further examination of the activities of amylase, lipase and lecithinase of *V. alginolyticus* cultured with different salinity (10‰, 20‰ and 30‰) showed no significant difference either (Fig. 2B). To further determine the effect of hyposalinity on the virulence of *V. alginolyticus* in infected oysters, we injected *V. alginolyticus* cultured with different levels of salinity (10‰, 20‰ and 30‰) into oysters and observed the mortality (Fig. 2C). The oyster death was first observed on one day post injection. Cumulative mortality of oysters at the end of the experiment was not significantly different among the three groups of different salinity (10‰, 20‰ and 30‰). Together, these results showed that changes in salinity level did not have a significant effect on growth and virulence of *V. alginolyticus*. Therefore, the effect of hyposalinity on the oyster’s mortality (Fig. 1) may not be caused by affecting *V. alginolyticus*, but the oysters.

**Fig. 2.**
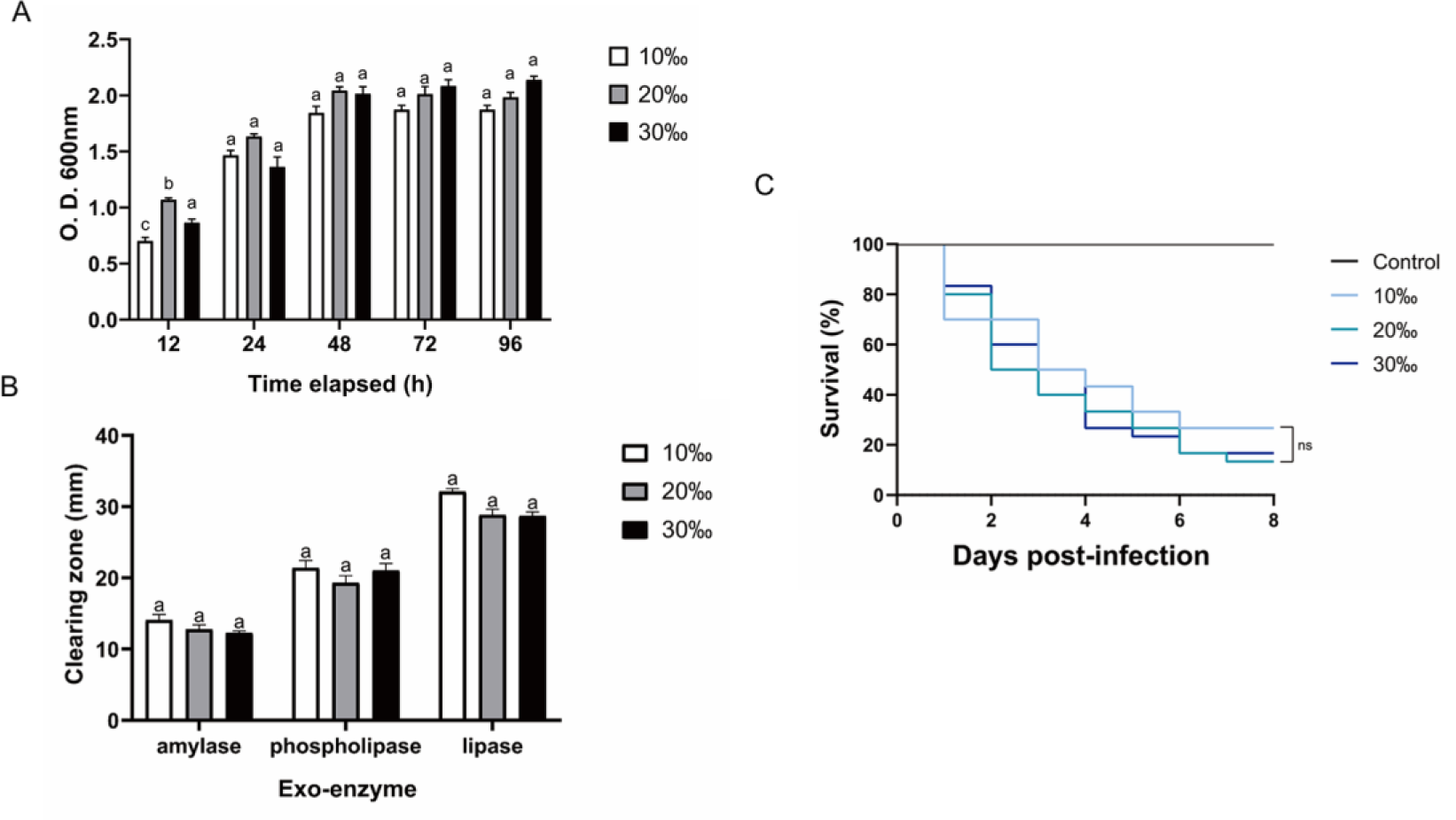
Effect of hyposalinity on growth and virulence of *V. alginolyticus*. (A) Effect of salinity on growth of *V. alginolyticus*. Bacterial amounts were determined at 12, 24, 48, 72 and 96 h. Each bar represents the mean (±S.E.) of three determinations. Bars for the same time with different letters are different at statistical significance (P < 0.05). (B) Effect of salinity on amylase, lipase and lecithinase activities of *V. alginolyticus*. Each bar represents the mean (±S.E.) of three determinations. Bars with different letters are significantly different (P < 0.05). (C) Survival analysis of oysters infected with *V. alginolyticus* cultured with different levels of salinity. n=90, in triplicate; ns: not significant.

### Impact of hyposalinity on digestive bacterial microbiota of the oysters

To investigate the microbiota dynamics of oysters infected with *V. alginolyticus* under normal salinity (denoted as “30‰Vibrio” hereafter) or hyposalinity (denoted as “10‰Vibrio” hereafter), we analyzed digestive bacterial microbiota using 16S rRNA sequencing. Overall, we obtained 1,518,721 raw sequencing reads from 24 samples (2 salinity gradients, 4 sampling times, in triplicate, Table S1). All the sequences were delineated into OTUs with 97% sequence similarity values. A sufficient sequencing depth was obtained as confirmed by species richness rarefaction curves (Fig. S1). Clearly, changes in microbiota composition were much higher in “10‰Vibrio” oysters than in “30‰Vibrio” oysters throughout the experiment (Fig. 3A). There were 27 genera with relative abundance higher than 0.1% in at least one sample, and the remaining genera were grouped as “Others”. The Chao1 and Shannon’s H indexes of alpha diversity increased significantly in “10‰Vibrio” oysters in comparison with “30‰Vibrio” oysters (Chao1: analysis of variance (ANOVA), df = 22; P = 0.010 and Shannon’s H index: ANOVA, df = 22; P =0.001) (Fig. 3B). Consistently, the principal coordinate analysis (PCoA) of the Bray-Curtis dissimilarity matrix (beta diversity) revealed a higher microbiota dispersion in “10‰Vibrio” oysters than in “30‰Vibrio” oysters (F = 2.330, R^2^ = 0.505; P = 0.035; PERMANOVA) (Fig. 3C). The hierarchical clustering heat map analysis at the genus level also showed that the microbial community structure of “10‰Vibrio” and “30‰Vibrio” was significantly different during infection of *V. alginolyticus* (Fig. 3D). Apparently, compared with “30‰Vibrio” oysters, bacteria associated with marine organism diseases including *Vibrio*, *Acinetobacter*, *Bacteroides* and *Streptococcus* became dominant genus in “10‰Vibrio” oysters during the process of infection.

**Fig. 3.**
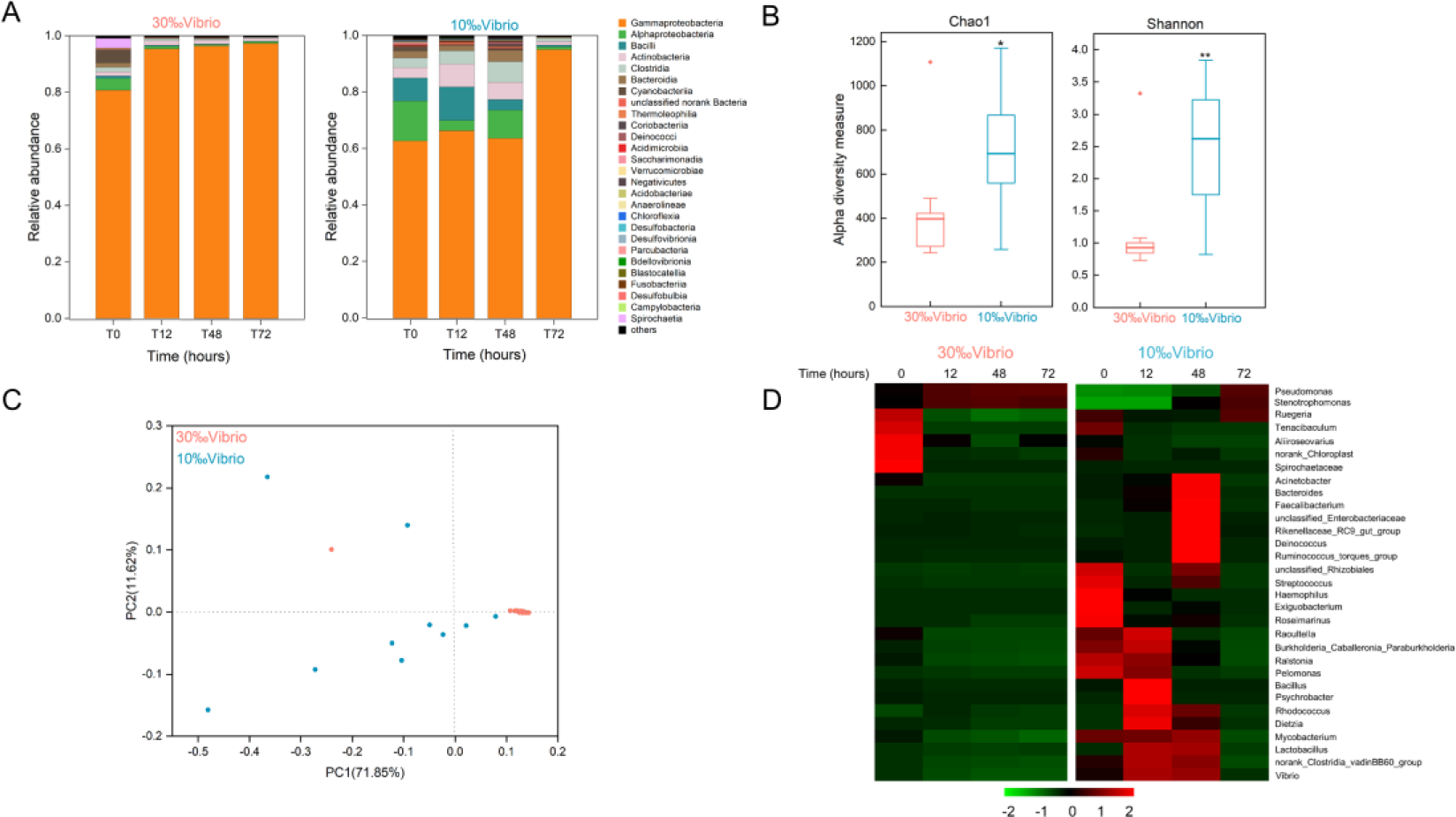
Microflora changes in *V. alginolyticus* infected oysters exposed to salinity of 10‰ and 30‰. **(A)** Relative proportion of bacteria (class level) for “10‰Vibrio” and “30‰Vibrio” oysters. T0, T12, T48 and T72 indicated different sampling time-points (in hours) during the experiment. **(B)** Temporal dynamics of alpha diversity in *V. alginolyticus* infected oysters at different salinity (10‰ and 30‰). Chao1 and Shannon’s H index for “30‰Vibrio” and “10‰Vibrio” oysters. Asterisk indicated P < 0.05. **(C)** Principal coordinate analysis (PCoA) plot of the Bray-Curtis dissimilarity matrix of the microflora comparing “10‰Vibrio” and “30‰Vibrio” oysters. The axes of the PCoA represent the two synthetic variables that explained the most variation of the dataset. **(D)** The abundance of each bacteria at the genus level in “10‰Vibrio” and “30‰Vibrio” oysters. Only genera with a relative proportion >0.5% in at least one sample are shown. The abundance of each type of bacteria in “10‰Vibrio” and “30‰Vibrio” oysters at the genus level during the *V. alginolyticus* infection. Green colour (smaller Log10 index) indicated lower abundance. Red colour (larger Log10 index) represented higher abundance.

### Impact of hyposalinity on immune response of the oysters

To determine how hyposalinity affects the host immune response to *V. alginolyticus*, we compared transcriptional profiles of oysters infected with *V. alginolyticus* at hyposalinity and normal salinity using RNA-seq. In total, total RNA of gill tissues collected from 0h, 12h and 48h (denoted as T0, T12 and T48, respectively) post infection at different salinity (10‰ and 30‰) were sequenced (Table S2), with 73.9-82.5% of reads being mapped to the *C. gigas* reference genome. Verification by RT-qPCR confirmed the results of the mRNA sequencing (Fig. S2; Table S5). Differentially expressed genes (DEGs) between each time point and the T0 control (“30‰Vibrio” versus T0, “10‰Vibrio” versus T0 at 12h and 48h post infection), and between the two groups (“30‰Vibrio” versus “10‰Vibrio” at 12h and 48h post infection) were identified (Fig. 4A). Comparison between each salinity group and T0 (“30‰Vibrio” versus T0 and “10‰Vibrio” versus T0) showed that the number of DEGs in infected oysters at hyposalinity was higher than that at normal salinity (Fig. 4A), suggesting that hyposalinity stress posed additional transcriptome alternations in addition with *V. alginolyticus* infection. To further clarify the specific factors that cause phenotypic differences in survival post infection under hyposalinity, we compared the two groups (“30‰Vibrio” versus “10‰Vibrio”), and identified a total of 3038 and 3187 DEGs in “10‰Vibrio” oysters compared with “30‰Vibrio” oysters at 12h and 48h post infection, respectively (Fig. 4A). To infer the biological processes regulated at hyposalinity, we performed gene ontology (GO) enrichment analysis (P < 0.05) (Table S3). Compared with the transcriptome of “30‰Vibrio” oysters, more functional categories related to immunity were enriched at “10‰Vibrio” oysters (26% at 12h; 27.8% at 48h). Interestingly, the vast majority of functional categories related to immunity were associated with inflammatory cytokines and apoptosis or cell death functional pathways (55% at 12h; 35.2% at 48h). Most of the genes involved in these pathways were induced in “10‰Vibrio” oysters (Fig. 4B; Table S3), such as “necrotic cell death”, “regulation of interleukin-1 beta production”, “interleukin-4-mediated signaling pathway”, “regulatory T cell differentiation”, and “intrinsic apoptotic signaling pathway in response to oxidative stress”. KEGG enrichment analysis of DEGs enabled further identification of significantly enriched pathways (P < 0.05) (Table S4). Compared with “30‰Vibrio” oysters, the “NF-Kappa B signaling pathway” and “TNF signaling pathway” were enriched in “10‰Vibrio” oysters at 12h and 48h post infection. In addition, the “apoptosis”, “necroptosis”, “Nod-like receptor signaling pathway”, “IL-17 signaling pathway”, “toll-like receptor signaling pathway” and “TH17 cell differentiation” were also enriched at 12h post infection. Taken together, we posited that the infected oysters had severely inflammatory response at low salinity compared with normal salinity, which were further verified by selection of representative genes. Sixteen DEGs related to cell death (apoptosis or necrotic death) and inflammation were chosen to verify the expression pattern between “30‰Vibrio” and “10‰Vibrio” (Fig. 4C; Table S5). The results showed that genes related to cell death (apoptosis or necrotic death) (*BIRC3*, *TRAF2*, *CYLD*, *SIRT6*), regulation of interleukin-1 beta production (*BIRC2*), and interleukin-4-mediated signaling pathway (*PARP14*), IL-17 signaling pathway (*PTGER4*), interferon-gamma-mediated signaling pathway (*IFI30*), TNF signaling pathway (*TNFaip3*), regulation of T cell differentiation (*NFKBIA*) and oxidative stress (*HSP70B2*, *HSPA12A*, and *HSPA13*) were expressed at higher levels in “10‰Vibrio” oysters. Consistently, expression of *BCL2L1* gene, which inhibits cell death, was suppressed in “10‰Vibrio” oysters. In summary, these results suggest that hyposalinity stress induced more intense inflammatory and cell death (apoptosis or necrotic death) responses in *V. alginolyticus* infected oysters.

**Fig. 4.**
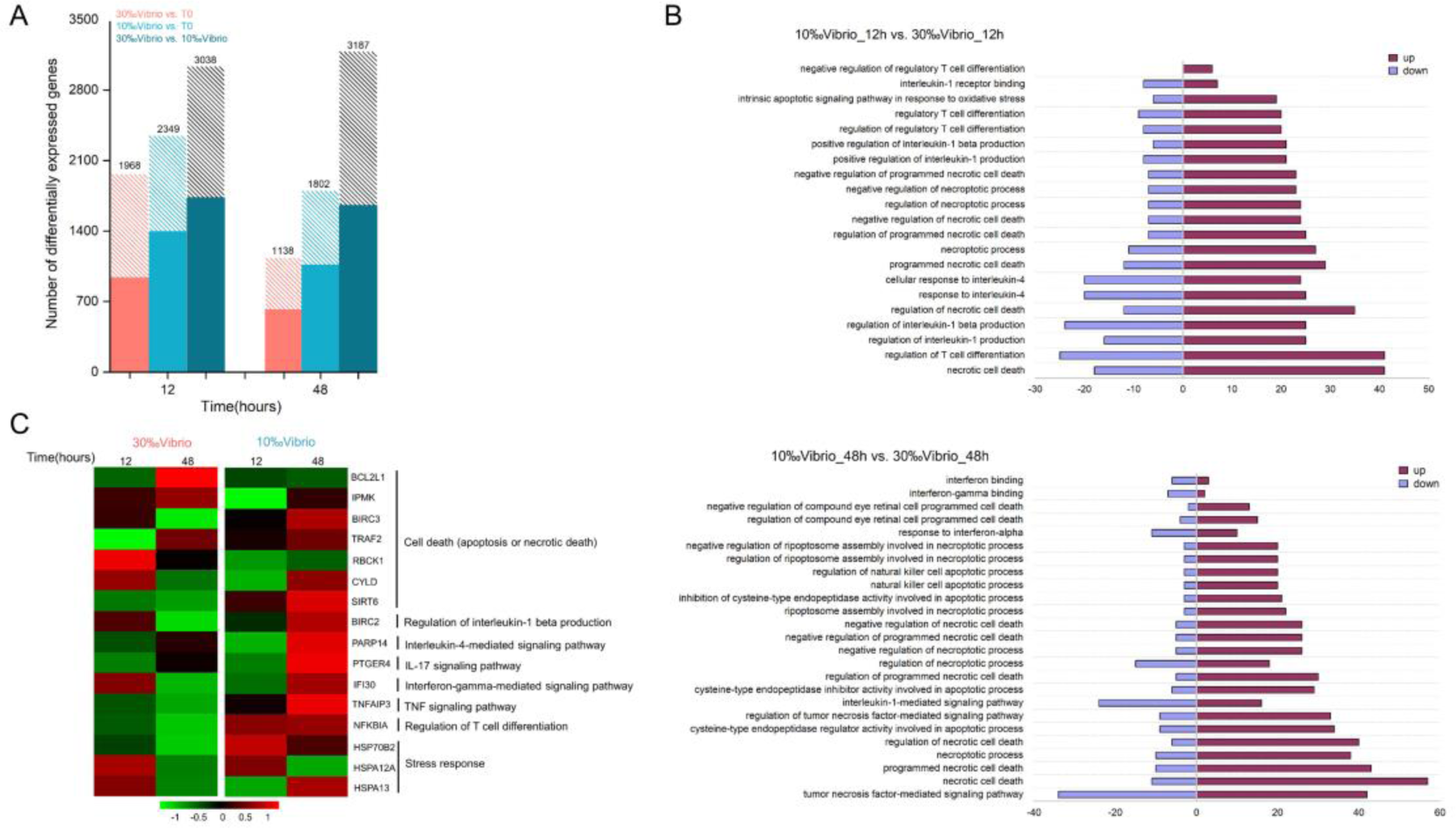
Comparative transcriptome profiling of *V. alginolyticus* infected oysters exposed to salinity of 10‰ and 30‰. **(A)** Identification of differentially expressed genes between *V. alginolyticus* infected oysters exposed to normal salinity (30‰) and hyposalinity (10‰) at 12h and 48h post infection. The filled colored bars indicated number of genes that were expressed at higher levels, while hashed bars indicated number of genes expressed at lower levels. **(B)** GO categories associated with necrotic cell apoptosis or death and inflammation that were significantly enriched at 12h and 48h post infection. The red-colored bars indicated genes were expressed at higher levels in 10‰Vibrio, while purple-colored bars indicated number of genes expressed at higher levels in 30‰Vibrio. **(C)** Verification of expression profiles of genes associated with necrotic cell apoptosis or death and inflammatory response. Differential expression analysis was performed between each time point and time zero. The intensity of the color from green to red indicates the magnitude of differential expression in log_2_(foldchange).

**Fig. 5.**
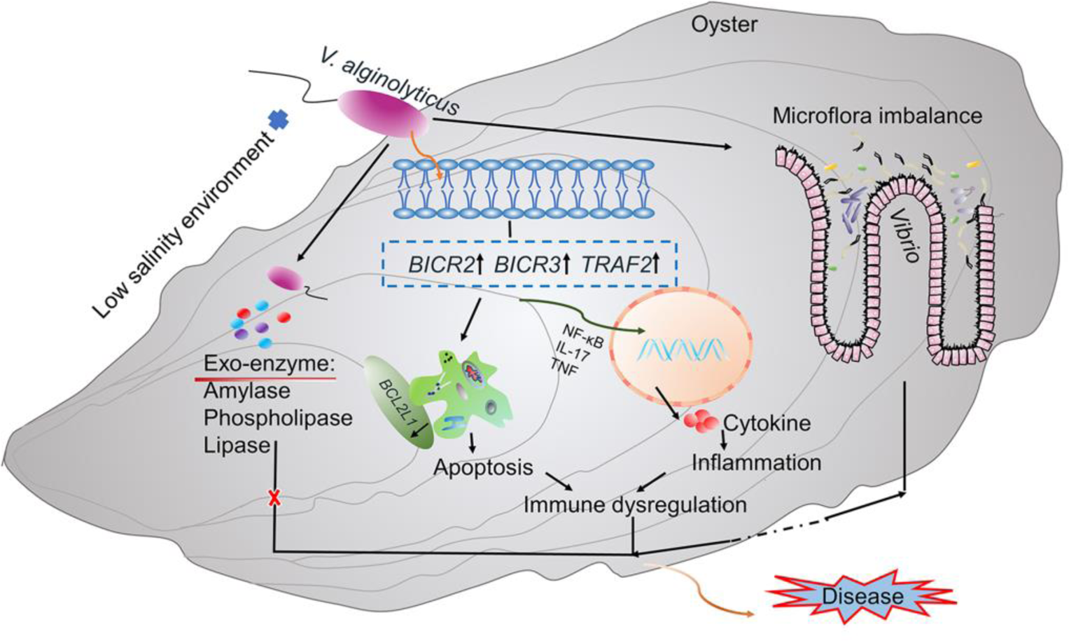
Microflora imbalance and immune dysregulation are hypothesized to cause high mortality of *V. alginolyticus* infected oysters exposed to low salinity environment. Low-salinity environment causes changes in digestive bacterial microbiota of infected oysters, leading to increased abundance of pathogenic bacteria such as *Vibrio* and disruption of microflora homeostasis of the host. Moreover, the low-salinity stress induced alterations of expression of genes involved in apoptosis and inflammation (e.g., *BICR2*, *BICR3*, *TRAF2*), which promotes production of inflammation-related cytokines that lead to immune dysregulation in oysters. Together, microflora imbalance and immune dysregulation caused by effect of hyposalinity stress drive high mortality of the oysters.

## DISCUSSION

Understanding of the interactions among environment factors, host genetics, and pathogens is crucial to unravel the complex pathogenic mechanisms in the host organism. In the present study, we observed the high mortality of oysters infected with *V. alginolyticus* at hyposalinity, and performed further investigation on their synergistic effect on the mortality of the infected oysters. The mortality rate of infected oysters exposed to hyposalinity (10‰) was 100%, in contrast that the mortality rate of infected oysters exposed to normal salinity (30‰) was 33%. The results showed that hyposalinity stress could trigger an increased mortality in oysters infected with pathogens, which is consistent with previous studies that reported a sharp increase in mortality of oysters exposed to extreme precipitation events (47). We further investigated the cause for mass mortality of infected oysters at hyposalinity (“10‰Vibrio”) and speculated whether hyposalinity stress has an impact on pathogen infection or host physiology, or both.

Changes in the natural environment affect the growth of pathogens and their production of toxins. Before the host’s immune defense system fully functions, fast-growing pathogens may secrete more virulence factors to destroy the host’s non-specific defense and lead to occurrence of disease (48). However, in the present study, the change of salinity had no significant effect on the growth and virulence of *V. alginolyticus*. This suggested that high mortality rate observed in “10‰Vibrio” oysters may not be caused by affecting growth and virulence of *V. alginolyticus*.

To better understand the relationship between the high mortality of infected oysters at hyposalinity and intestinal microbiota composition, bacterial communities in the digestive gland of oysters were examined. Our results showed that alterations in microbiota composition and diversity in “10‰Vibrio” oysters were more significant than that in “30‰Vibrio” oysters. Previous studies have reported that salinity stress and pathogen exposure can cause changes in intestinal microbial composition of various aquatic organisms (49–51), while the occurrence of many diseases is associated with changes in intestinal microbial composition (52). Intestinal microorganism can provide protection against pathogens by producing inhibitory compounds and competing for nutrients and space. Therefore, we hypothesized that such changes in the intestinal microbiome caused by hyposalinity might exacerbate disease progression in infected oysters.

The finding of *Gammaproteobacteria* as the dominant class of intestinal microbiota in infected oysters is consistent with reports in previous studies (28,49,53). Hyposalinity stress affected the relative abundance of *Gammaproteobacteria* in the oyster bacterial community. The OTUs with increased abundance in “10‰Vibrio” oysters were assigned to the genera *Vibrio*, *Acinetobacter*, *Bacteroides*, and *Streptococcus*, all of which have been previously associated with diseases in some aquatic organisms. There is growing evidence of the role of *Vibrio* communities in the development of oyster disease (54–60). Specifically, previous studies revealed that native *Vibrio* community was replaced by pathogenic *Vibrio* species prior to oyster disease onset (61), especially when oysters were exposed to stressors that may facilitate the transition to more pathogen-dominant communities (28, 58). Our work also suggested the increased abundance of *Vibrio* community when infected oysters were exposed to hyposalinity. Although the relative abundance was low, it can have a significant influence on the host health (62). In addition, *Acinetobacter* (63), *Bacteroides* (64–66), and *Streptococcus* (67–69) were over-represented in “10‰Vibrio” oysters, which have been identified as known pathogens in fish or crab. Hyposalinity stress may disrupt the homeostasis of the microbial composition in infected oysters, likely leading to proliferation of various opportunistic pathogens and inflammation in oysters. The stable microbiome of “30‰Vibrio” may suggest their potential role in host adaptation to stressors (70–73).

Comparative expression profiling of Vibrio-infected oysters at low and normal salinity provided insights into molecular basis associated with high mortality of oysters under biotic and abiotic stresses. Functional analysis of DEGs revealed that gene pathways related to necrotic cell apoptosis or death, interleukin-1 production, interleukin-4 mediated signaling pathway, and T cell differentiation were significantly enriched in “10‰Vibrio” oysters. Interestingly, most of the genes involved in these significantly enriched functional pathways were upregulated upon hyposalinity stress. It has been reported that necrotic cell apoptosis or death is a critical determinant for the initiation of inflammation, and damage-associated molecular patter (DAMP) signals sent by dead cells can attract more monocytes, inducing a vicious cycle of inflammation and accelerating disease development (74). Interleukin-1 (IL-1) family cytokines IL-1β play key roles in inflammation (75), whose excessive and/or dysregulated activity can cause common inflammatory disorders (76, 77). Naive CD4^+^ T cells can be differentiated into Th1 cells that produce IFN-γ and TNF-α, Th2 cells that produce IL-4, and Th17 cells that produce IL-17 during intracellular bacterial infection (78, 79). The excessive cytokines may induce an imbalance in the Th1/Th2 ratio, thereby enhancing immune activation and inflammatory response, which is closely related to the onset and severity of colitis, inflammatory bowel disease (IBD) and asthma (80).

Inflammation protects the host from pathogens and can repair damaged tissues. However, excessive inflammation can cause tissue damage and malaise (81). We further validated expression profiles of critical genes involved in cell death (apoptosis or necrotic death), including *BIRC3*, *TRAF2*, *CYLD*, and *SIRT6*, inflammatory factors, including *BIRC2*, *PARP14*, *PTGER4*, *IFI30*, *TNAFAIP3*, and *NFKBIA*, and oxidative stress including *HSP70B2*, *HSPA12A*, and *HSPA13*. We showed that all of these genes were expressed at higher levels in infected oysters exposed to hyposalinity stress. These results were consistent with the results of high expression of apoptosis and inflammatory cytokine related genes in immune-damaged fish and chickens (82–85). In addition, cell death and inflammation can interact to form an auto-amplification loop that causes organ damage (44).

Functional analysis of DEGs also showed that some immune functions (i.e. defense response to bacteria, complement activity, antimicrobial peptide biosynthesis, antigen processing and presentation) were significantly enriched in infected oysters at hyposalinity. Nevertheless, as shown by the high mortality of infected oysters at low salinity, an increase in immune factors does not necessarily indicate higher resistance to microbial disease. These results suggest that low salinity stress induces more severe inflammatory and apoptotic responses in infected oysters, and these factors may interact to cause immune damage and disrupt basal immune homeostasis, similarly as the phenomena that have been reported in fish (45, 86). Hyposalinity creates unfavorable environment for effective immune defense against pathogen infection in oysters. However, mechanism by which hyposalinity induces immune damage in infected oysters deserves further investigations.

It is well known that healthy intestinal flora is closely related to maintaining body health by regulating host immune homeostasis (87). In our study, the increase in abundance of pathogenic bacteria corresponded to the increased inflammatory response in “10‰Vibrio” oysters. Therefore, we speculated that, on one hand, low salinity may indirectly affect immune homeostasis by altering the oyster’s digestive bacterial microbiota homeostasis in infected oysters; on the other hand, hyposalinity may indirectly affect host intestinal microbiota homeostasis by inducing immune changes in infected oysters. At present, in-depth studies on the complex interactions between oyster intestinal microbiome and host immunity are lacking. Future work focusing on this process warrants better understanding of its role in surveilling host health and pathogenesis.

## CONCLUSIONS

In this study, we performed an experiment by mimicking a sudden drop of salinity and pathogen infection with *V. alginolyticus* to investigate their synergistic effect on the mortality of the infected oysters toward understanding of the interaction among environment, host, and pathogen. The infected oysters exhibited higher mortality rate under hyposalinity stress compared to normal salinity. Further investigation revealed that high mortality rate was not due to effect of hyposalinity stress on promoting growth and virulence of *V. alginolyticus*, but likely due to disruption of homeostasis of digestive bacterial microbiota in infected oysters, leading to the bursts of pathogenic bacteria as well as excessive inflammatory response to cause immune dysregulation. The interaction of these factors ultimately impairs the oyster’s capacity to defend against infection of *V. alginolyticus* at low salinity, causing mass mortality. This work provides significant information in decoding mechanisms of synergistic interaction among environment factors, host genetics, and digestive microbiota, and how they contribute to pathogenesis.

## MATERIALS AND METHODS

### Experiment animals

Healthy oysters (Pacific oyster, *Crassostrea gigas*, wet weight of 18.2±2.8 g) obtained from an oyster farm (salinity of 30‰) in Rongcheng (Shandong, China) were used for experiment. The oysters were transported to laboratory and acclimatized in 30‰ seawater for two weeks. Continuous aeration was provided and the water quality was monitored. The oysters were fed with concentrated *Chlorella vulgaris* ad libitum. The water quality parameters during experimental challenge trials are follows: pH at 8.1-8.2, dissolved oxygen at 8.0-9.0ppm, salinity at 30-32, and nitrite at 1-3ppm. Unused feed and faecal matter were siphoned out daily and 25% water was changed every other day.

### Hyposalinity stress and *V. alginolyticus* infection

Oysters were randomly assigned to glass tanks containing 20L seawater. The pathogenic *V. alginolyticus* strain isolated and purified previously (15) was inoculated in 50mL 2216E broth. The 5×10^8^ CFU mL^-1^ bacterial suspension was used for infection. The oysters were anesthetized with magnesium chloride (MgCl_2_, 50g/L) solution before being injected with *V. alginolyticus*. Each oyster was intramuscular injected with 50 μl (5×10^8^ CFU/oyster) bacterial suspension volume through a microsyringe (100±0.5μl). After injection, infected oysters were placed in 20L glass tanks containing 10‰ (“10‰Vibrio”) and 30‰ (“30‰Vibrio”) seawater, respectively (n = 30 / glass tank, in triplicate). Low salinity seawater is prepared using 24h aerated tap water and seawater prepared through natural sand filtration. Oysters injected with an equal volume of artificial seawater and cultured in 30‰ seawater were used as unchallenged control (“Control”). During the experiment, digestive gland tissues of oysters were sampled at 0, 12, 48 and 72h post infection for bacterial flora analysis. Gill tissues were sampled at 0, 12, and 48 h post infection for transcriptomic analysis. During sampling, nine individuals were randomly selected at each time point, and the three gills or digestive glands were combined into one sample (three replicates) and stored in liquid nitrogen.

### Impact of hyposalinity on growth and virulence of *V. alginolyticus*

The *V. alginolyticus* was cultured in 2216E broth at 28℃ for examination of growth and virulence. *V. alginolyticus* was inoculated in 50 mL flask containing 10 mL tryptone soy broth in different concentrations of sterile saline (10‰, 20‰ and 30‰) for bacterial growth test at 28±1°C. Each test was conducted in triplicate and bacterial growth was monitored at 12, 24, 48, 72, and 96h of incubation by measuring the optical density (OD) at 600 nm.

For the determination of extracellular enzyme activity of *V. alginolyticus*, overnight cultures of the bacterial strain in tryptone soy broth media in different concentrations of sterile saline (10‰, 20‰ and 30‰) were diluted to an OD 600 of 0.5. Except for the lipase assay, 10 μl of the diluted cultures was spotted in the middle of the test plates. For the lipase test, the strain was serially diluted and spread-plated on top of the lipase test agar. All assays are done in triplicate. In this assays, 2216E agar plates supplemented with 0.5% starch, 1% Tween 80, or 1% egg yolk emulsion (88), respectively, were used. The development of colorless and transparent circle around the colonies was observed for amylase after dripping Lugol’s iodine solution. The development of opalescent zones around the colonies was observed for lipase and phospholipase. The diameter of the zones was measured after 2-4 days of incubation at 28℃.

The experiment for the determination of virulence of *V. alginolyticus* in oysters, *V. alginolyticus* was cultured in tryptone soy broth media in different concentrations of sterile saline (10‰, 20‰ and 30‰) for 24 hours, and then *V. alginolyticus* was harvested by centrifugation at 8000×g for 5 min at 4°C. The pellet was re-suspended in saline solution (0.85% NaCl) at 5×10^8^ CFU mL^-1^ as the stock bacterial suspension for injection. Ten oysters were used for each of the treatment (10‰, 20‰) and control groups (30‰). The experimental infection was done in triplicate. Each oyster was injected with a total of 50μl of bacterial suspension into the adductor muscle. After injection, oysters were placed in 20L glass tanks containing 30‰ seawater, and observed for one week. During the experiment, water temperature was maintained at 20±1°C. Control oysters were injected with an equal volume of sterile saline solution.

### Bacterial microbiota analysis

The total genomic DNA was isolated from digestive gland samples using the E.Z.N.A.^®^ soil DNA kit (Omega Bio-tek, Norcross, GA, USA) according to the manufacturer’s instructions. The genomic DNA quality was confirmed by running 1% agarose gel electrophoresis. PCR was performed using two universal bacterial primers 341F (5’-CCTAYGGGRBGCASCAG-3’) and 806R (5’-GGACTACNNGGGTATCTAAT-3’). The PCR assays were carried out in triplicate as follows: 95 °C for 3 min, followed by 27 cycles of 95 °C for 30 s, 55 °C for 30 s, 72 °C for 45 s, and a final extension of 72 °C for 10 min. All PCR products were extracted from 2% agarose gel, and purified with a AxyPrep DNA Gel Extraction Kit (Axygen Biosciences, Union City, CA, USA). The DNA concentration of each PCR product was determined using a QuantiFluor™-ST fluorescent quantitative system (Promega, USA), and mixed with the appropriate proportion based on sequencing requirements. Sequencing was performed using an Illumina MiSeq platform (Illumina, San Diego, USA) for 300bp paired end reads.

Sequences were demultiplexed and quality-filtered using QIIME (version 1.9.1). Operational taxonomic units (OTUs) were clustered with 97% similarity cutoff using UPARSE (version 7.1) and chimeric sequences were identified and removed using UCHIME (89). The taxonomy of each 16S rRNA gene sequence was analyzed by RDP Classifier (version 2.2) against the SILVA (version 138) 16S rRNA database using confidence threshold of 70% (90). OTUs that reached 97% similarity were used for alpha diversity estimations, which included diversity (Shannon), richness (Chao I), and rarefaction curve analysis using Mothur (Version 1.30.2). Significant differences of alpha diversity index among groups were calculated with one-way ANOVA. PCoA was conducted according to the Bray-Curtis distance matrix calculated using OTU information from each sample. A heat map was computed using relative abundances and the gplot package of R software. Differences among populations were analyzed using a one-way ANOVA. P < 0.05 was considered statistically significant.

### Transcriptome profiling of infected oysters

The gill samples of two salinity groups (“10‰Vibrio” and “30‰Vibrio”) were used for transcriptome analysis. Total RNA was extracted from with Trizol Reagent (Invitrogen) according to the manufacturer’s instructions. The RNA quality was confirmed by running 1% agarose gel electrophoresis. RNA concentration and purity were measured using NanoDrop (Thermo Fisher Scientific), and the RNA integrity number (RIN) was assessed using the RNA Nano 6000 Assay Kit of the Bioanalyzer 2100 system (Agilent Technologies). The kit used for library construction was the NEBNext^®^ Ultra^™^ RNA Library Prep Kit for Illumina^®^ (NEB, USA). Total RNA extracted from the samples was subject to mRNA enrichment using poly-T oligo-attached magnetic beads. The extracted mRNA was randomly broken by NEB Fragmentation Buffer into short fragments. The fragmented mRNA was used as template for synthesis of first-strand cDNA using random hexamer primer and M-MuLV Reverse Transcriptase (Rnase H-). The second-strand cDNA was synthesized using DNA polymerase I and RNase H. After purification, end repair, adenylation of 3’ ends of DNA fragments, and adaptor ligation, cDNA fragments of 250-300 bp were selected using AMPure XP beads and enriched by PCR. After the libraries were tested to be qualified, the paired-end sequencing was performed using Illumina HiSeq 2500 platform.

Raw reads in fastq format generated from Illumina sequencing were assessed by FastQC. Clean reads were obtained by trimming reads containing adapter, reads containing poly-N, and reads with low sequencing quality. The downstream analyses were based on the high-quality clean reads. The reference oyster genome (GCA_902806645.1) was first indexed (91), and then the paired-end clean reads were aligned to the indexed reference genome using Hisat2 (v2.2.1) (92). The counts of reads mapped to each gene were obtained using FeatureCounts (v1.6.0) (93). The fragments per Kilobase of transcript per million mapped reads (FPKM) of each gene was then determined based on the length of the gene and counts of reads mapped to the gene (94). DEseq2 (1.30.1) was used for differential gene expression analysis. A model based on the negative binomial distribution was used to calculate the P-value. The resulting P-values were adjusted using the Benjamini and Hochberg approach for controlling the false discovery rate. In order to obtain significantly differentially expressed genes (DEGs), the criteria were set as P- value < 0.05 and |fold-change| > 1. GO functional analysis and KEGG pathway analysis of the DEGs were performed to identify enriched gene pathways.

### Quantitative Real-Time PCR

We selected 12 differentially expressed genes for quantitative real-time PCR (qRT-PCR) analysis. Specific primers for qRT-PCR were designed according to the reference sequences using Primer Premier 5.0 (Table S5). The EF-1α gene was used to detect baseline expression of mRNA in the gill tissues of different salinity groups. The amplification was performed on the LightCycler 480 real-time PCR instrument (Roche Diagnostics, Burgess Hill, UK) using SYBR^®^ Premix Ex Taq^™^ (TaKaRa). The reaction conditions were as follows: 95°C for 5 minutes; 95°C for 5 s, 58°C for 30 s and 72°C for 30 s, 40 cycles in total. The solubility curve of PCR products was performed to ensure specific amplification. Relative gene expression levels were calculated by the 2^−ΔΔCt^ method (95). Data were analyzed by t-test using software SPSS 24.0, and P-value < 0.05 was considered as statistical significance.

### Data availability

All genomic and transcriptomic sequence data used in this study have been deposited in the Sequence Read Archive (SRA) of National Center for Biotechnology Information with the BioProject accession number of PRJNA756403 and PRJNA756710.

## ACKNOWLEDGEMENTS

This work was supported by the grants from National Natural Science Foundation of China (No. 31802293 and No. 41976098), the Young Talent Program of Ocean University of China (No. 201812013), and the Fundamental Research Funds for the Central Universities (No.202042011).

## AUTHOR CONTRIBUTIONS

S.L. conceived the study and obtained the funding. X.L. and S.L. designed the experiment. X.L., H.W. performed the experiment, X.L., B.Y. and C.S. analyzed the data. X.L. drafted the manuscript, S.L. revised the manuscript. Q.L. supervised the work. All authors have read and approved the final manuscript.

## CONFLICT OF INTEREST

The authors declare no conflict of interest.

